# Monocarboxylate Transport in *Drosophila* Larval Brain during Low and High Neuronal Activity

**DOI:** 10.1101/610196

**Authors:** Andres Gonzalez-Gutierrez, Andrés Ibacache, Andrés Esparza, L. Felipe Barros, Jimena Sierralta

## Abstract

The transport of lactate and pyruvate between glial cells and neurons plays an important role in the nervous system metabolic coupling. However, the mechanisms and characteristics that underlie the transport of monocarboxylates (MC-T) *in vivo* are poorly described. Here we use *Drosophila* expressing genetically-encoded FRET sensors to provide an *ex vivo* characterization of the MC-T in motor neurons and glial cells from the ventral nerve cord. We show that lactate/pyruvate transport on glial cells is coupled to protons and is more efficient than in neurons. Glial cells maintain higher levels of intracellular lactate generating a positive gradient towards neurons. Moreover, our results show that under increased activity lactate and pyruvate rise on motor neurons and suggest that this depends on the transfer of lactate from glial cells mediated in part by the previously described MC transporter Chaski, giving support to the *in vivo* glia to neurons lactate shuttling during activity.

## INTRODUCTION

A hallmark of the brain is its high energy requirement^1^. This energy demand is fulfilled by a constant supply of glucose and oxygen, which are mostly used to fuel the recovery of ion gradients challenged by synaptic transmission and action potentials^2,3^. Other energy-rich molecules that may be used to fuel brain cells are the monocarboxylates (MC) L-lactate, pyruvate and ketone bodies^4–7^. *In vitro* studies show that neurons and glial cells are able to take up these substrates despite displaying markedly different metabolism, with glial cells being less dependent on oxygen than neurons. Whether the intercellular transport of MC is relevant *in vivo* and which are the cells in the mammalian brain that produce and used them is still a matter of intense debate and research^8–13^.

Invertebrates such as *Drosophila* and *C.elegans* display a brain organization different from the one in mammals, maintaining characteristics such as the presence of glial cells as well as a blood brain barrier. Recent evidence supports that similarly to vertebrate organisms there are metabolic differences between neurons and glial cells, suggesting that some fundamental features of brain metabolism are conserved. It was shown that *Drosophila* glial cells *in vitro* release alanine and lactate when fueled by trehalose or glucose and that knockdown of glycolytic genes in glia, but not in neurons, leads to severe neurodegeneration^14^. Moreover, it was suggested that *Drosophila* glial cells provides lactate to neurons, which use it as a substrate to form fatty acids that in turn are transported back to glial cells where are stored as lipid droplets^15^. Despite the increasing evidence of the role of lactate, there is little knowledge about the characteristics of the movement of lactate between glial cells and neurons, particularly during activity.

In mammalian cells, the transport of lactate and other monocarboxylates is mediated by proton-linked monocarboxylate transporters (MCTs, the SLC16A family)^16–18^. These electroneutral transporters move one molecule of lactate along with a proton using both gradients but also can function as an exchanger of monocarboxylates. In *Drosophila*, there are fifteen genes with a putative role as MCT (*d*MCT) and just a few of them have been studied. The gene CG8271, called silnoon (*sln*), was identified as a *d*MCT and shown to be involved in apoptosis induced by extracellular butyrate^19^. Our group identified Chaski (CG3409) as a *d*MCT for lactate/pyruvate^20^. In the brain, *chaski* plays an important role in synaptic activity, locomotion and in the adaptive response to nutritional restriction; the latter being rescued by the expression of *chaski* in glial cells. In order to better understand the brain metabolism in *Drosophila* it is necessary to determine the main features of lactate transport, both in basal and in activity conditions as well as the transporters involved.

Genetically-encoded fluorescent biosensors allow the study of metabolism in specific cell types with high temporal resolution. Using such sensors in *Drosophila*, it was shown that glucose is fast and freely distributed between glial and neuronal cells by transporters not yet identified^21^. In addition, another study showed that an increase in energy flux into the neurons of mushroom body in the *Drosophila* adult brain (the center associated with memory and learning in insects) precedes and drives long-term memory formation^22^. The source and nature of the molecules driving the increased energy consumption could not be established due to the lack of basic information about the fluxes of energy-rich molecules in the *Drosophila* brain.

Here we provide an *ex vivo* characterization of monocarboxylate transport in glial cells and motor neurons from *Drosophila* larval ventral nerve cord. Using FRET-based genetically encoded sensors for metabolites, we were able to detect transport of lactate and pyruvate in all subtypes of glial cells studied as well as in motor neurons. Our experiments show that the transport of monocarboxylate is faster in glial cells than in neurons while the basal level of lactate is lower in neurons than in glia, evidencing a standing gradient from glial cells to neurons. We determine that the transport of lactate and pyruvate is coupled to the transport of protons and that *d*MCTs undergo substrate-induced trans-acceleration, as their mammalian counterparts. Furthermore, we found that neuronal activity triggers fast increase in intracellular lactate in astrocyte-like glial cells, which is followed by a slower increase in neuronal lactate. Using *chaski* mutants we provide evidence of the role of Chaski in glia but importantly give a strong support to the transport of lactate from glia to neurons during neuronal activity. Together these results not only show that *d*MCTs share key features of vertebrates MCTs but confirms the role of *chaski* mediated lactate transport in glial cells giving support to the notion of activity-dependent glia to neuron lactate shuttling.

## RESULTS

### Lactate and pyruvate transport in motor neurons and glial cells

To study the transport of lactate and pyruvate in *Drosophila* larval neurons and glial cells, we conducted experiments in an *ex vivo* preparation in which isolated brains from third instar larvae were perfused with a saline solution containing trehalose, the main carbohydrate present in larval hemolymph^14,23^. The pH of freshly isolated larval hemolymph was 6,7± 0,06 (N=20), consistent with previous reports^24^. Thus, in our experiments the saline solution pH was adjusted to 6.7. Using the Gal4/UAS system^25^, we expressed the FRET-based sensors for lactate and pyruvate *Laconic* and *Pyronic*^26,27^ in motor neurons using the specific driver *OK6-Gal4*^28^ or in glial cells using the driver *repo-Gal4* (Figure 1A). Brains were exposed to increasing lactate concentrations from 1 mM, near to the levels found in the hemolymph (747 ± 74 µM, n=4). Lactate up to 1 mM induced an increase in lactate both in motor neurons (*OK6Gal4>Laconic*) and glial cells (*repoGal4>Laconic*, Figure 1B-C). Although the increase in lactate at 1 mM lactate seems to occur at similar rate in both type of cells, higher concentrations led to higher lactate in neurons.

**Figure 1.**
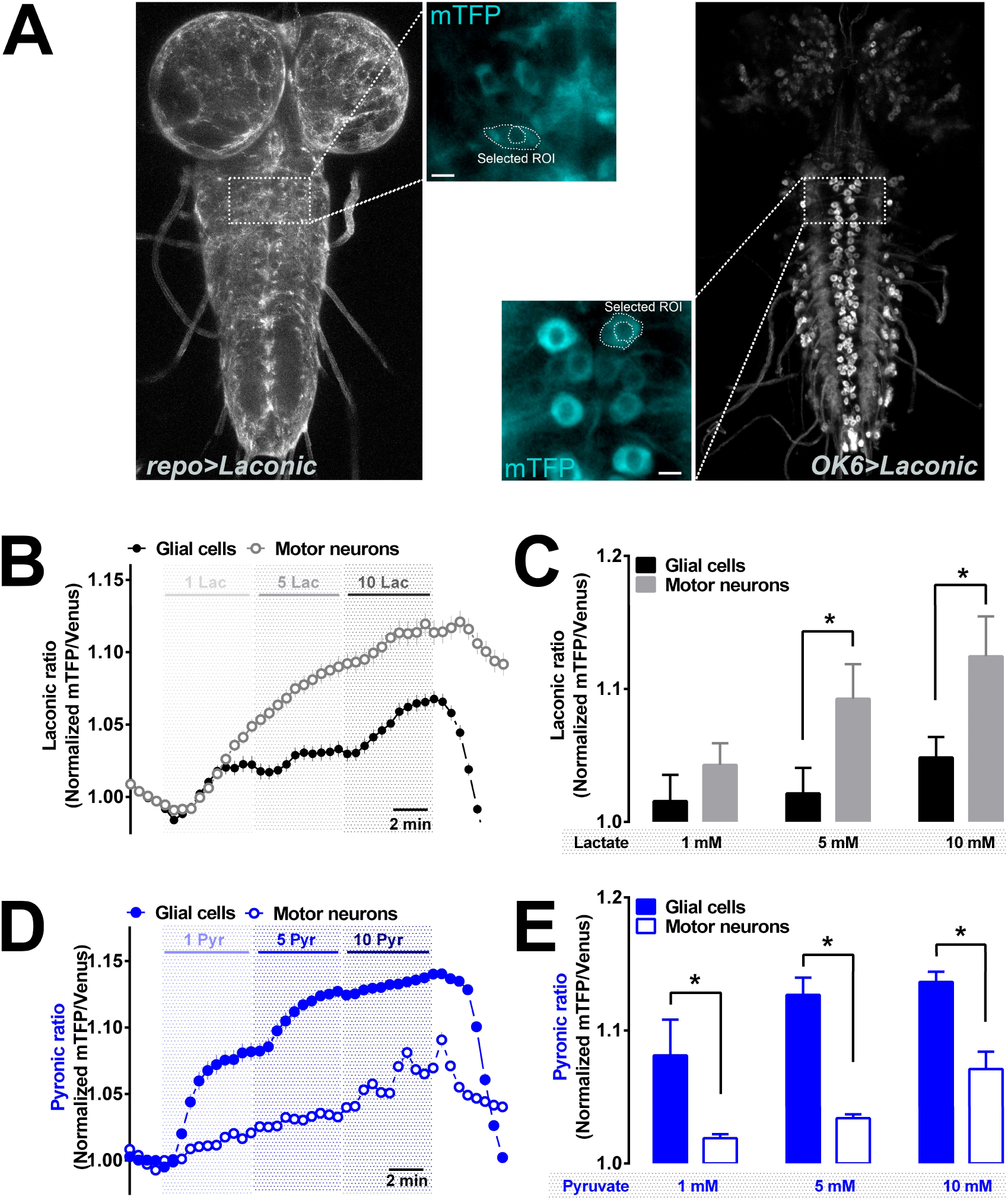
Lactate and pyruvate transport in motor neurons and glial cells from *Drosophila* larvae. **(A)** Confocal images of third instar larvae brains expressing *Laconic* in glial cells (*repoGal4>Laconic, left*) or motor neurons (*OK6Gal4>Laconic, right*). Images show mTFP fluorescence from *Laconic* sensor (excitation: 438 nm; emission: 480 nm). Zoom from the images placed in the middle shows the procedure to select regions of interest analyzed. Scale bars represent 5 μm. **(B)** Recordings of *Laconic* signals from glia cells (*repoGal4>Laconic*, black circles) or motor neurons (*OK6Gal4>Laconic*, grey open circles) brains exposed to increasing lactate concentrations (1mM, 5 mM and 10 mM lactate). Images were taken every 30 seconds. **(C)** Mean fluorescence ratio of mTFP/Venus obtained from the last 2 minutes of each stimulus from over 60 cells from 6 independent experiments. Each symbol represents mean ± SE. *p<0.05. **(D)** Recordings of *Pyronic* signals from brains expressing this sensor in glial cells (*repoGal4>Pyronic*, blue closed circles) or in motor neurons (*OK6Gal4>Pyronic*, blue open circles) exposed to increasing pyruvate concentrations (1mM, 5 mM and 10 mM pyruvate). **(E)** Mean fluorescence ratio of mTFP/Venus obtained from the last 2 minutes of each stimulus from over 50 cells from 5 independent experiments. *p<0.05.

Exposing larval brains to increasing pyruvate concentrations resulted in accumulation of pyruvate within both types of brain cells. However, contrary to lactate, the response of pyruvate was larger in glial cells. (Figure 1D-E). In fact, the response elicited in neurons by exposure to 10 mM pyruvate was equivalent to that observed in glial cells in response to 1 mM pyruvate, suggesting that glial cells display a higher gradient for pyruvate than neurons and/or higher *d*MCTs density. Interestingly, we observed that after washing out lactate or pyruvate, the return of *Laconic/Pyronic* signal to basal levels occurred faster in glial cells (for *Laconic*: −0.043±0.008 min^−1^, N=6; for *Pyronic*: −0.044±0.006 min^−1^, N=5) than motor neurons (for *Laconic*: −0.010±0.002 min^−1^, N=6; for *Pyronic*: −0.010±0.001 min^−1^, N=5). In summary, using these genetic-encoded FRET sensors we were able to measure both lactate and pyruvate fluxes in glial cells and motor neurons and show that the transport of these MCs is markedly different among these cells in the *Drosophila* larval ventral cord.

### pH dependence and trans-acceleration of *Drosophila* MCTs

Mammalian monocarboxylate transporters show two distinctive functional features, proton-coupling and trans-acceleration (also termed accelerated-exchange). The coupling to protons makes the transport sensitive to the pH gradient^29^. To investigate whether *d*MCTs are also proton-coupled we measured intracellular pH using the genetically-encoded sensor *pHerry*^30^. *pHerry* combines the pH-sensitive green fluorescent protein Super-Ecliptic pHluorin (SE-pH)^31^ and the red fluorescent protein mCherry, used to normalize expression levels and pH-independent changes^32^. We expressed *pHerry* in motor neurons (*OK6Gal4>pHerry*) or in glial cells (*repoGal4>pHerry*) and determined changes in *pherry* fluorescence after exposing the brain to different lactate/pyruvate concentrations. Both glial cells and neurons showed a transient acidification after lactate exposure, consistent with proton influx (Figure 2A). The pH changes were faster and larger in glial cells than in neurons. Exposure of the brain to pyruvate elicited significant changes in glial pH (Figure 2B). The acidification followed a time course similar to that of pyruvate (Figure 1D) suggesting proton-coupled pyruvate uptake. In motor neurons, exposure to pyruvate caused small changes in *pHerry* signal in agreement to the small changes observed in pyruvate (Figure 1D). Changes in intracellular pH due to lactate/pyruvate transport in glial cells closely resembled the transport of these molecules measured by *Laconic* and *Pyronic* sensors indicating that lactate/pyruvate transport is coupled to H^+^ transport. On the other hand, pH changes in motor neurons do not completely correlated with the lactate/pyruvate changes measured using the FRET sensors. This suggests that lactate/pyruvate transport in motor neurons is less coupled to protons or that stronger pH buffering/extrusion mechanisms in motor neurons muffled the acidification. Another non-exclusive explanation is that pyruvate, which is mainly captured by glia, has limited access to motor neurons in this brain preparation.

**Figure 2.**
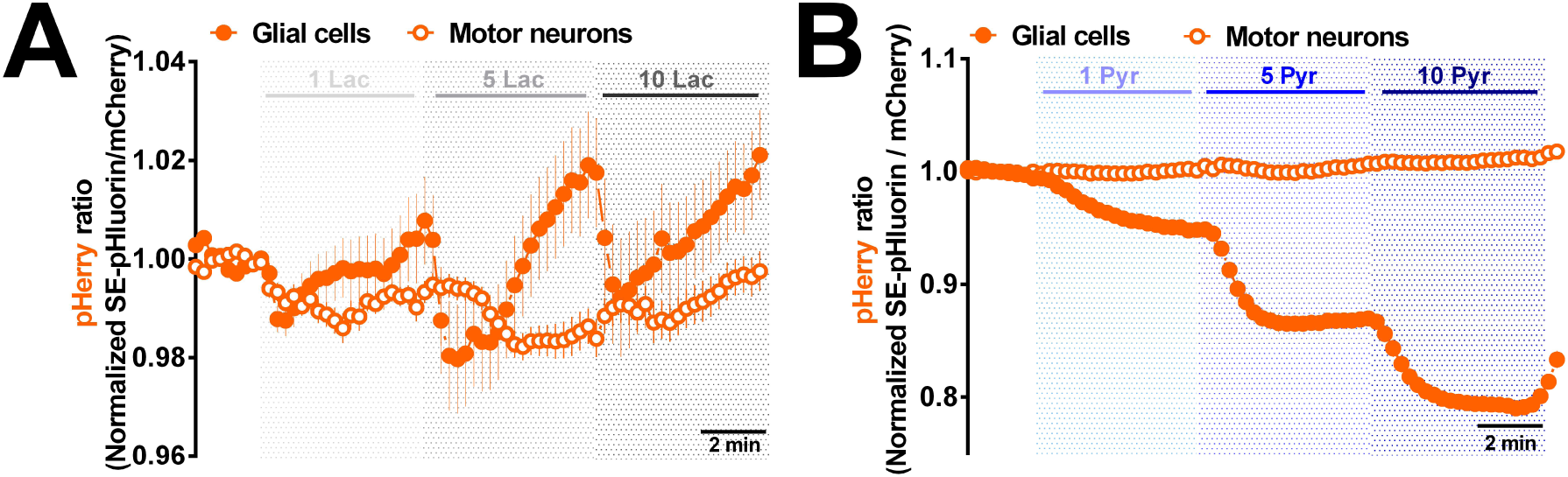
Lactate and pyruvate induces changes on *pherry* signals in motor neurons and glial cells. **(A)** Recordings from brains expressing the pH sensor *pHerry* in glial cells (*repoGal4>pHerry*, orange closed circles) or in motor neurons (*OK6Gal4>pHerry*, orange open circles) exposed to increasing lactate concentrations (1mM, 5 mM and 10 mM lactate). Images were taken each 15 seconds. Traces are the mean fluorescence from *pHerry* ratios obtained from 4 independent experiments (over 50 cells) for each Gal4 used. **(B)** Using the same protocol, we recorded brains expressing *pHerry* in glial cells (*repoGal4>pHerry*, orange closed circles) or in motor neurons (*OK6Gal4>pHerry*, orange open circles) but exposed to increasing pyruvate concentrations (1mM, 5 mM and 10 mM pyruvate). Images were taken each 15 seconds. Traces are the mean fluorescence from *pHerry* ratios obtained from 4 independent experiments (over 50 cells) for each Gal4 used.

The second feature showed by mammalian MCTs is trans-acceleration, which is the increased transport of a given monocarboxylate (e.g. lactate) induced by the presence of a monocarboxylate at the opposite site of the membrane (e.g. pyruvate)^33^. We explored this property in *Drosophila* brains by exposing the organ to 10 mM pyruvate. As shown in Figure 3A, the increase in pyruvate was accompanied by a decrease in lactate. Although the lactate decrease is demonstrative of trans-acceleration, its time course was much slower than the increase in pyruvate, suggesting that the lactate pool is being replenished by pyruvate conversion by LDH. Neurons responded with increases in both pyruvate and lactate (Figure 3B). As in these experiments the brain is intact and neurons are surrounded by glial cells, our explanation for these effects is that pyruvate is first converted into lactate within glial cells, and then shuttled into neurons. To test these hypotheses, we repeated the trans-acceleration experiment using the non-metabolized pyruvate analog oxamate^20^. Oxamate caused a faster depletion of glial lactate than pyruvate (pyruvate: −0,003 ± 0,002 min^−1^; oxamate: −0,008 ± 0,001 min^−1^) but did not affect neuronal lactate (Figure 3C). This result supports the idea that the neuronal lactate increase observed in pyruvate-exposed brains was caused by glial lactate production from pyruvate and that in these conditions neuronal lactate levels are low. As a low lactate concentration in basal condition could preclude the observation of trans-acceleration in neurons, we increased tissue lactate by exposing larval brains to high lactate in the presence of ammonium chloride to block mitochondria and stimulate glycolysis^34,35^. Then, we repeated the trans-acceleration experiment with oxamate. Under these conditions (high intracellular lactate), a clear trans-acceleration was could be induced by oxamate exposure in motor neurons (Figure 3D). Taken together, these results demonstrate that under basal conditions neuronal lactate is kept low and that lactate generated in glial cells may shuttle to neurons.

**Figure 3.**
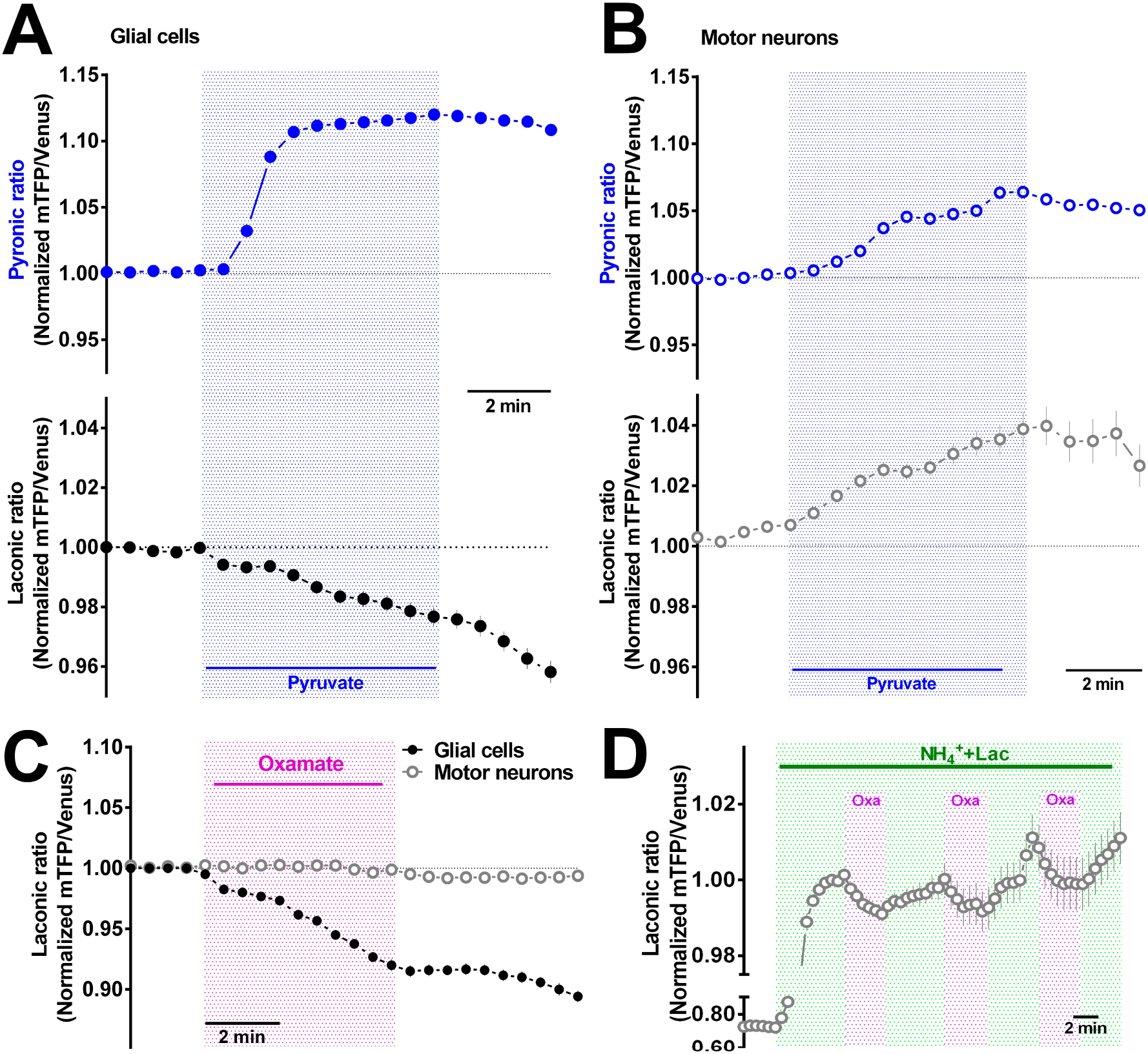
*Drosophila* monocarboxylate transporters undergo trans-acceleration in *ex vivo* preparation. **(A)** Brains expressing in glial cells the pyruvate sensor *Pyronic* (*repoGal4>pyronic*, blue closed circles in upper panel) and the lactate sensor *Laconic* (*repoGal4>Laconic*, black closed circles at the bottom) were exposed to 10 mM of pyruvate. Traces are mean fluorescence from 4 independent experiments (40 cells) for *Pyronic* and 6 independent experiments for *Laconic* (71 cells). **(B)** The same experiment was performed in brains expressing *Pyronic* in motor neurons (*OK6Gal4>pyronic*, blue open circles in upper panel) and the lactate sensor *Laconic* (*OK6Gal4>Laconic*, black circles at the bottom) and exposed to 10 mM of pyruvate. Traces are mean fluorescence from 4 independent experiments for *Pyronic* (40 cells) and 5 independent experiments for *Laconic* (56 cells). **(C)** Brains expressing *Laconic* in glial cells or motor neurons (*repoGal4>Laconic*, black closed circles, *OK6Gal4>Laconic*, grey open circles, respectively) were exposed to 10 mM of the non-metabolizable pyruvate analog oxamate. Values are mean fluorescence from 6 independent experiments for glial cells (71 cells) and 5 independent experiments for motor neurons (56 cells). **(D)** Brains expressing *Laconic* in motor neurons (*OK6Gal4>Laconic*, grey open circles) were constantly bathed with 10 mM NH_4_Cl + 10 mM lactate and then exposed to 10 mM of oxamate. Values are mean signal of *Laconic* from a representative recording (12 cells).

Using the ability of *d*MCTs to undergo trans-acceleration, we determined whether there is a difference on lactate basal levels between neurons and glia cells as our previous experiments suggest and as it has been seen in vertebrates^35^. We incubated brains with 1 mM lactate to load the cells with levels of lactate in equilibrium with the extracellular solution until the *Laconic* signal was steady. Then, we induced the extrusion of lactate changing the lactate by oxamate and compared the change in the *Laconic* signals between glia and neurons. Supplementary figure 1 shows that *Laconic* signal decreased faster in glial cells compared to neurons (−0.045 min^−1^ vs −0.010 min^−1^) and in higher degree (−13.5 ± 2.3% vs −2.2 ± 0.3%) in response to oxamate, showing that glial cells maintain higher concentrations of lactate (approx. 5 times more) than neurons.

Taken together, these results show that *d*MCTs share the key features with their vertebrate counterparts which are co-transport of protons and trans-acceleration. They also show that MC transport is more efficient in glial cells than in neurons and that in basal conditions, glial cells maintain higher concentrations of lactate than neurons, which results in lesser lactate gradient and trans-acceleration easier to induce.

### Monocarboxylate transport in subtypes of glial cells

The first glial cells in direct contact with hemolymph which acts as a filter for solute transport into the brain are the perineurial an subperineurial glial cells (PG and SPG). Then, solutes can be transported to internal glial cells as cortex-glia (CG) that surround neuronal bodies, astrocyte-like glia (AG), surrounding the neuropile and ensheathing glia (EG) covering axons (Figure 4A). Unpublished data from our group indicates that all glial cells express several genes predicted to be *d*MCTs, including *chaski* ^20^. Thus, given the diversity of subtypes of glia we wanted to determine if other types of glia also show lactate transport. For this we repeated the experiment of exposure to increased concentrations of lactate using specific drivers for subtypes of glia. Figure 4B shows that all subtypes of glial cells studied increased *Laconic* signals in response to a rise in extracellular lactate, although not all glia responded to 1 mM lactate. Interestingly, in SPG cells (*R54C07Gal4>Laconic*) *Laconic* signal rises significantly and fast after being exposed to 1 mM lactate. This increase in the signal is transient, a phenomenon that is not observed in the other types of glia, which maintain the signal until the next concentration of lactate with minor attenuations. This suggests that SPG maintain low levels of lactate providing a gradient for the influx of lactate, which is then efficiently transported out from SPG to internal cells and interstitium. Notably, CG (*R55B12Gal4>Laconic*) exhibited a strong increase in *Laconic* signal at 5 mM lactate exposure compared to the rest of glial cells studied, suggesting that cortex glial cells may differ in the level of expression of *d*MCTs and/or the basal intracellular lactate content. Surprisingly, in ensheathing glial cells (*R83E12Gal4>Laconic*) we only see increases in *Laconic* signal at 5 and 10 mM of extracellular lactate concentrations. This could be explained by the sequestration or metabolization of lactate by other glial cells, being these types of cells less exposed to lactate. Taken together, these results show that in basal conditions most glial cells show a reduced lactate transport comparing to motor neurons, suggesting higher basal levels of lactate (smaller gradient for the transport) similar to what was reported in mammalians.

**Figure 4.**
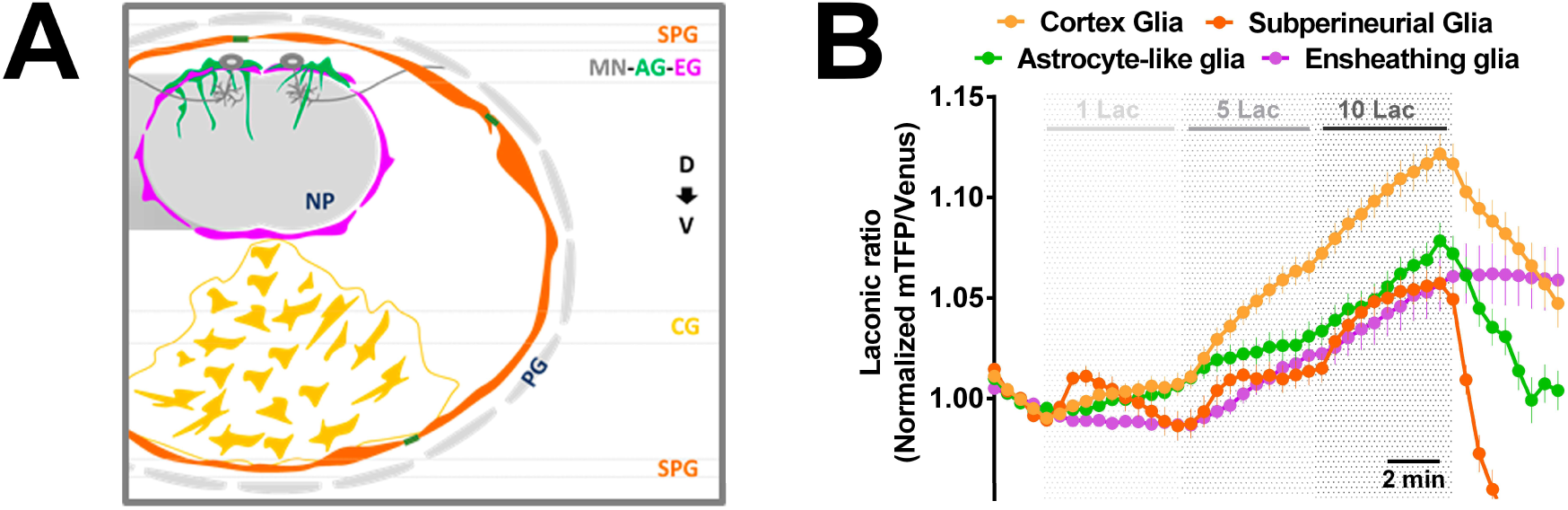
Monocarboxylate transport in subtypes of glial cells. **(A)** Schematic figure of a cross-section of ventral nerve cord showing the distribution of glial cells and dorsal motor neurons in the ventral nerve cord of *Drosophila* (PG: perineurial glia; SPG: subperineurial glial cells; CG: cortex glia; AG: astrocyte-like glial cells; EG: ensheathing glia; MN: motor neuron; D: dorsal VNC; V: ventral). **(B)** Brains expressing *Laconic* in subperineurial glial cells (*R54C07Gal4>Laconic*), cortex glia (*R55B12Gal4>Laconic*), astrocyte-like glia (*AlrmGal4>Laconic*) and ensheathing glia (*R83E12Gal4>Laconic*) were exposed to increasing lactate concentrations (1mM, 5 mM and 10 mM lactate). Values are mean fluorescence ratio of mTFP/Venus obtained from over 50 cells from 4-5 independent experiments for each Gal4 line used.

### Activity-induced lactate changes in glial cells and neurons of *Drosophila*

From previous studies in mammalian cells, it has been proposed that glial-derived lactate can supply the energy requirements for neuronal function. It is still a matter of debate whether this mechanism operates *in vivo*, during resting states or during activity^11,36^, as neuronal glycolysis is also activated during neurotransmission^37,38^. In *Drosophila*, glial glycolysis is necessary for survival, suggesting that transport of lactate from glial cells fuels neurons^14^. To explore the activity-induced transfer of monocarboxylates between glia and neurons, we determined lactate transport in motor neurons and glial cells after inducing an increase in the neuronal activity by adding the GABA_A_ receptor antagonist Picrotoxin (PTX)^39,40^. To confirm the increase in activity, larval brains expressing the genetic-encoded calcium sensor *GCaMP6f* in motor neurons or in astrocyte-like glial cells, were exposed to PTX (80 μM). Figure 5A shows that PTX at this concentration and with a delay of one minute after being applied can induce a large increase in *GCaMP6f* fluorescence in motor neurons which is followed by calcium spikes that occur with high frequency. *GCaMP6f* signals in AG do not show a significant change after PTX exposure. Under these conditions, in addition to lactate and pyruvate, we evaluated changes in intracellular glucose using the FRET-based sensor *FLII^12^Pglu700*µ*∆6*. As before the media contained only trehalose as source of energy. Additionally, we focused in AG glia (*Alrm-Gal4>UAS-FRET-sensors*).

**Figure 5.**
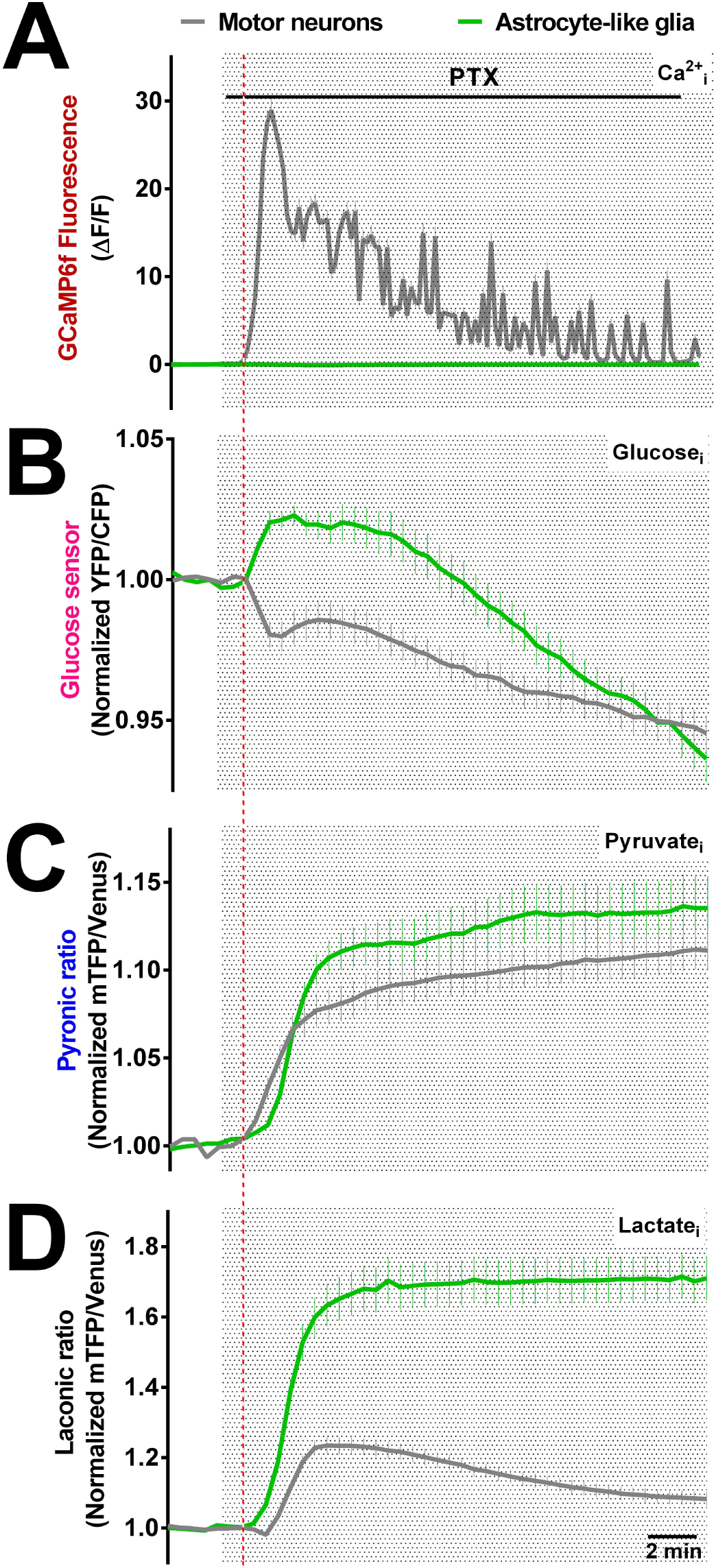
Picrotoxin-induced neuronal activity increased *Laconic* signal more in glial cells than motor neurons. Brains expressing the calcium sensor *GCaMP6f* **(A)**, the glucose sensor *FLII*^*12*^*Pglu700µ∆6* **(B)**, *Pyronic* **(C)** or *Laconic* **(D)** in astrocyte-like glial cells (*AlrmGal4, green lines*) and motor neurons (*OK6Gal4, grey lines*) were exposed to the GABA_A_ receptor antagonist Picrotoxin (PTX, 80 µM). For *GCaMP6f* fluorescence, correspond a representative recording for each Gal4 line used. For glucose, pyruvate and lactate sensors, values are mean fluorescence obtained from over 40 cells from 4 independent experiments for each Gal4 line used.

AG glia responded to evoked neuronal activity with a fast increase in lactate that reached a stable plateau (Figure 5D). Neurons displayed a lactate dip, followed by a transitory increase in lactate. Pyruvate on the other hand, both in neurons and glia showed a rapid rise followed by a secondary slower rise (Figure 5C). Glucose behaved differently in glia and neurons, in AG we observed a fast increase that was followed by a slower and sustained decrease that reached below control levels (Figure 5B). Neurons on the contrary responded with a fast decrease followed by a transient stabilization and a late decrease. These results show that neural activity causes rapid changes in the energy metabolism of brain cells in *Drosophila* larval brain and that PTX exposure provides a simple and reliable method to study brain metabolism under increased neuronal activity in this preparation.

### *Chaski* mutants exhibit a reduced transport of lactate in glial cells of *Drosophila*

Previously we characterized Chaski (Chk) as a *d*MCT with a role in synaptic activity in the NMJ of the larvae as well as locomotion and in the adaptive response to nutritional restriction in the adult *Drosophila*^20^. Unpublished data from our laboratory show that chk mutants are not able to survive mild starvation or stressful temperature during larvae stage, suggesting a more important role than previously thought. To determine the role of Chaski in this *ex vivo* system, wild type (W^1118^) or chk mutants (*chk*^*CRISPR*^) brains expressing *Laconic* in glial cells (*repo>Laconic*) or motor neurons (*C380>Laconic*) were exposed to increasing lactate concentrations. Figure 6A-B show that the response to lactate in glial cells was significantly reduced in *chk*^*CRISPR*^ comparing to control larvae. In motor neurons, the opposite was observed, with *Laconic* signal increased at high lactate concentrations in mutants compared to control larvae (Figure 6C-D). Thus, our results suggest that the absence of *chaski* leads to a decreased transport of lactate in glial cells and an upregulated lactate transport in motor neurons.

**Figure 6.**
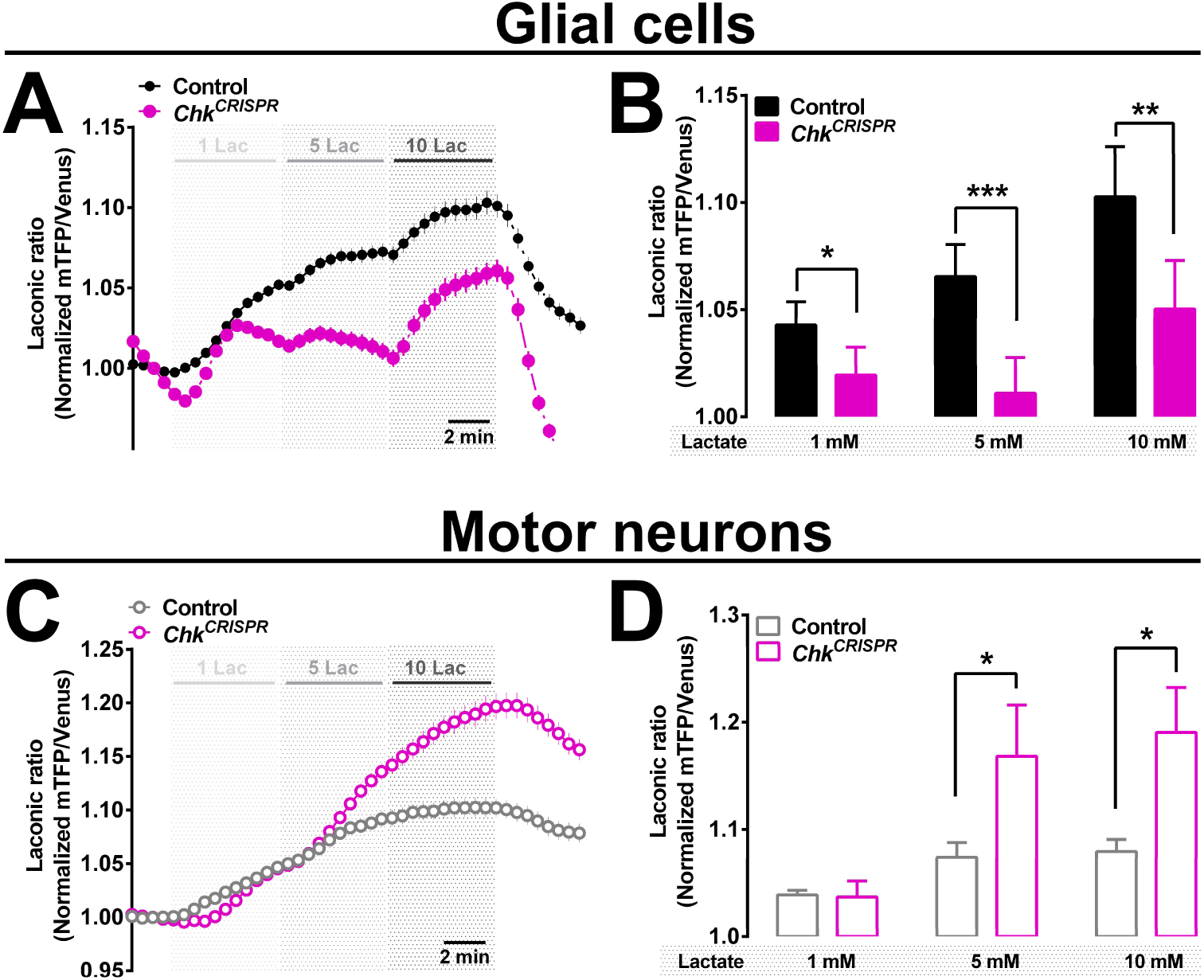
Reduced *Laconic* signal in glial cells from Chaski mutants exposed to lactate. Recordings of control larvae brains (W^1118^) and chaski mutant (Chk^CRISPR^) brains expressing *Laconic* in glial cells (*repoGal4>Laconic*, in **A**) or motor neurons (*C380Gal4>Laconic*, in C) and exposed to increasing lactate concentrations (1mM, 5 mM and 10 mM lactate). Values are mean fluorescence from 6 independent experiments (60 cells) for glial cells and 7 independent experiments (77 cells) for motor neurons. (**B** and **D**) Mean fluorescence ratio of *Laconic* fluorescence were obtained in the last two minutes of each lactate exposure. *: p<0.05; **: p<0.01; ***: p<0.001.

Using *chaski* mutants, we repeated the PTX experiments and measured lactate in neurons and glia cells. We aimed to determine whether the pyruvate rise in neurons induced by neuronal activity (Figure 5C) is a product of influx of lactate from AG cells. Figures 7A-B show that in *chk*^*CRISPR*^ flies the accumulation of lactate induced by neuronal activity in glial cells was substantially larger than in control flies. In contrast, the rise in lactate induced by neuronal activity was smaller in neurons (Fig. 7C-D). A parsimonious explanation for this mirroring behavior is that Chaski contributes to the transport of lactate between glial cells and neurons.

**Figure 7.**
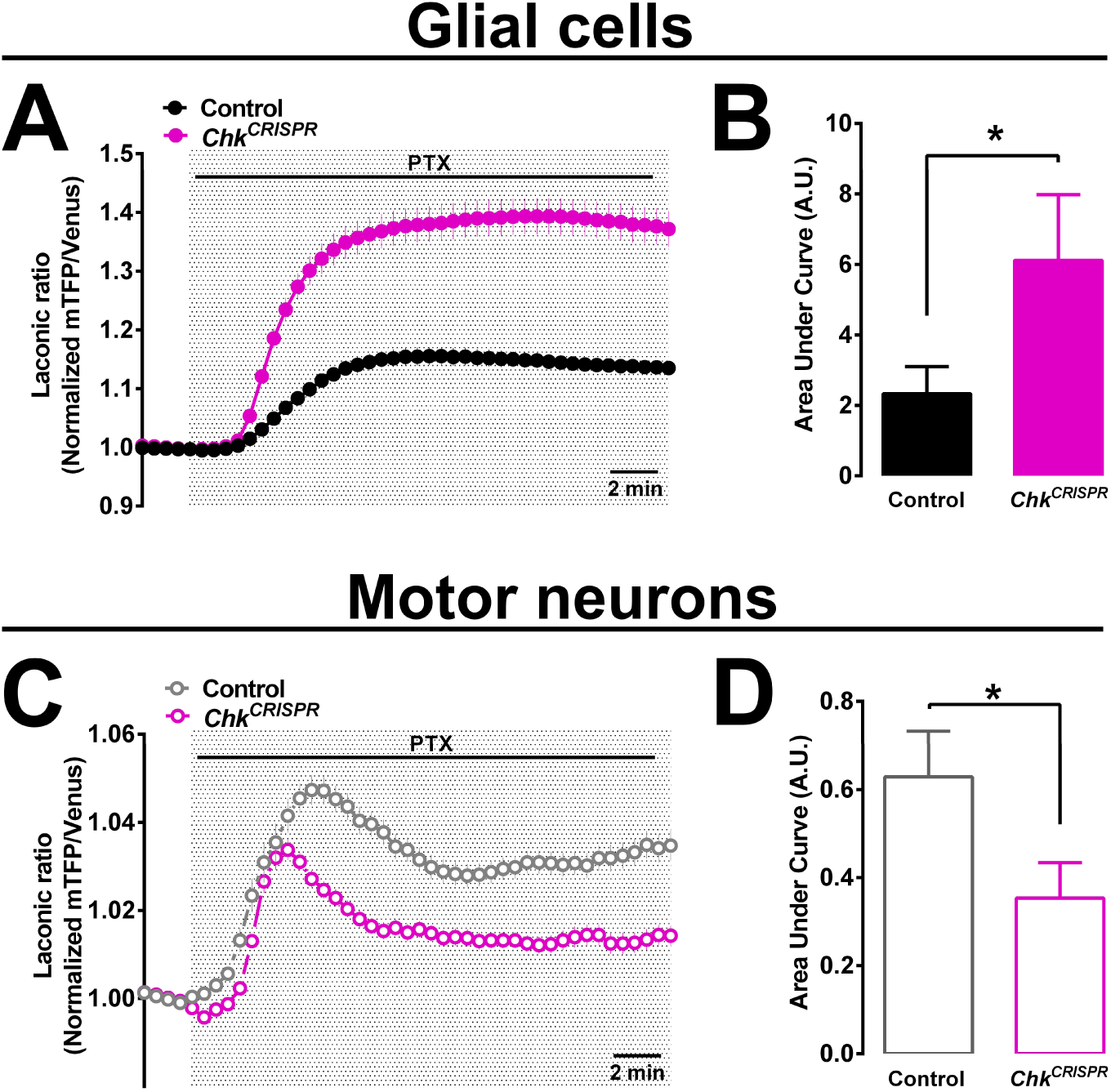
*Chaski* mutants shows altered response of *Laconic* signal after increased neuronal activity. Control larvae brains (W^1118^) and chaski mutant (Chk^CRISPR^) brains expressing *Laconic* in glial cells (*RepoGal4>Laconic*, in **A**) or motor neurons (*C380Gal4>Laconic*, in C) were recorded before and after being exposed to the GABA_A_ receptor antagonist Picrotoxin (PTX, 80 µM). (**B** and **D**) Intracellular lactate accumulation measured as the area under the curve of experiments in A and C. Values are mean ± SE from over 60 cells and 5 independent experiments.

## DISCUSSION

Recent evidence has shown the existence of a metabolic coupling between glial cells and neurons in the nervous system of *Drosophila*, where the production and transport of lactate and pyruvate plays an important role in synaptic activity, energy production and adaptation to nutritional stress^20^. However, the mechanisms and characteristics that underlie the transport of monocarboxylates *in vivo* have not been addressed.

### Main features of monocarboxylate transport between glial cells and neurons

Using genetically encoded FRET-based sensors we determined intracellular changes of lactate, pyruvate and glucose and were able to show *in situ* transport of lactate and pyruvate in glial cells and motor neurons. Using a genetically encoded pH sensor we also determined that monocarboxylate transport is coupled to the transport of protons, suggesting that hemolymph pH could modulate the degree of lactate and pyruvate transport inside the nervous system. It is known that hemolymph pH in *Drosophila* can be regulated by the nutritional state and even the type of food consumed^41^. Under our *Drosophila* culture conditions, we determined a pH near to 6.7 in freshly isolated hemolymph. Using a pH of 7.2, a pH more regularly used in *ex vivo* preparations of *Drosophila* larvae we could only detect lactate flux in motor neurons but not in glial cells, even at high extracellular lactate concentrations (Supplementary figure 2). This suggests that the lactate gradient between the hemolymph and the glia is small or that glial *d*MCT are more sensitive to pH than the ones present in neurons. Using the pH sensor and the lactate sensor we showed that glial cells display faster monocarboxylate transport than neurons (Figure 2A-B), a difference that may relate to the level of expression of *d*MCTs in each type of cells and/or the *d*MCT gene expressed by them. Mammalian neurons predominantly express MCT2, whereas astrocytes and oligodendrocytes express MCT1 and MCT4^42^. The lack of antibodies for *d*MCTs and a complete description of which candidate genes behaves as MCT and where they are expressed limits further characterization. However, we have already demonstrated that Chaski is mainly expressed in glia while Liu *et al* suggested silnoon as a neuronal MCT^15^.

Our data also show that *d*MCTs in neurons and glia undergo trans-acceleration, a characteristic of mammalian MCTs. Our experiments showing that the exposure to pyruvate to induce trans-acceleration resulted in pyruvate transport into the neuron and its fast conversion to lactate (Figure 3B) suggests that in these conditions lactate concentrations are higher in glial cells (low gradient to lactate influx) than in neurons (high gradient to lactate influx). We confirmed this hypothesis inducing the depletion of intracellular lactate in glial cells and neurons by exposing brains to oxamate after loading the cells with lactate. The comparison of basal levels of *Laconic* signal indicates that glial cells maintain higher lactate levels than neurons as in mammals at resting conditions ^35^. These results support the idea that glial cells from *Drosophila* acts as lactate producers, harboring a fast system of efflux that given the low intracellular levels in neurons favors the transport from glia to neurons.

*Drosophila* nervous system contains several subtypes of glia, each one having roles that mimics those played by astrocytes, oligodendrocytes or Schwann cells from vertebrates^43,44^. For example, SPG cells forms the blood-brain barrier and regulates the transport of small solutes into the nervous system covering not only the brain but also peripheral axons^45^; cortex-glia (CG) wraps neuronal cell bodies and is probably related with metabolic functions; astrocyte-like glia (AG), presents high functional and morphological similarities with vertebrates astrocytes and is in close contact with synapses in the neuropile, finally ensheathing glia (EG), surrounds axons. We could demonstrate that these glial cells differ in lactate transport capacity and kinetic (Figure 4B), highlighting a possible role of SPG as a mediator of fast transport of monocarboxylates inside the nervous system. Given their distribution and the fact that outer glial cells can generate free glucose from trehalose, the high rates of monocarboxylate transport that display subperineurial and cortex glial cells suggest that by these means neurons could be constantly fueled with energy-rich compounds.

### Production and transport of monocarboxylates under increased neuronal activity

There are emerging evidence pointing to an important role of monocarboxylate transport and metabolism in brain function and survival of flies^14,15,22^. However, there is still much debate about their essential role *in vivo*. In our work we found that the increased neuronal activity induced by picrotoxin leads to a high lactate and pyruvate production in astrocyte-like glia (AG, Figure 5C-D), cells that are in close contact with synapses in neuropile regions^43,46^. The origin of this production must come from the metabolization of glucose, since the extracellular media lacks lactate and pyruvate. Given the transient increase in glucose production in AG, the glucose source could be the breakdown of trehalose in outer glial cells or glial glycogen. Further studies will have to investigate which of this alternative is the main source. On the other hand, motor neurons show a moderate increase of lactate and relevant production of pyruvate after PTX-induced activity. Given the low glucose amount that the cells maintain and the very low glycogen deposits that have been described in neurons it is unlikely that neuronal glucose explains the important pyruvate production observed (Figure 5C). Our results and the literature are compatible with glial lactate been shuttle to motor neurons where is transformed to pyruvate for energy production by mitochondria. An active field of discussion has been how relevant can be glial-derived lactate as an energy substrate for neurons in vertebrate models^11,36^. Experiments conducted in mouse hippocampal slices indicates that electrically stimulated neurons use glucose for energy production, being lactate shuttle from glia to neurons dispensable^37^. However, in *Drosophila* the fact that the lack of key glycolytic enzymes in glial cells and not in neurons leads to neurodegeneration, strongly suggest that energy-rich metabolites derived from glial cells are crucial to sustain neuronal survival^14^. In our work we directly determined the role of the *d*MCT Chaski in the transport of lactate from glial cells to neurons. The absence of Chaski leads to an accumulation of lactate in glial cells during increased neuronal activity, due to reduced exit from cells (Figure 7A). Motor neurons that lacks Chaski showed an increased transport of lactate, probably due to an upregulation of another (or more than one) yet undetermined *d*MCT (Figure 6C). On these conditions, activity-induced intracellular lactate increase was reduced in motor neurons associated to a decreased efflux from glial cells (Figure 7C). If this influx of lactate is necessary for energy production in glutamatergic motor neurons is suggested by our previous report which indicates that *chk* mutants exhibit a locomotor phenotype that may be explained by an altered calcium handling due to a decreased energy production used to maintain ionic homeostasis in neurons^20^. Although the mechanisms underlying lactate shuttle from glia to neurons in *Drosophila* are only starting to be unveiled, our work shows that cellular and molecular components in the larval brain resembles vertebrate mechanisms^47^. For example, AG display properties of mammalian astrocytes that allow to study neuron-glia interactions^48^ including the expression of excitatory amino acid transporter 1 (*d*EAA1)^49–51^. Despite multiple similarities, the known differences should be incorporated in order to give a complete landscape of the metabolic interactions present in this system. Notably, glutamatergic presynaptic neurons acting on glutamatergic motor neurons in *Drosophila* are inhibitory and the excitatory presynaptic neurons are cholinergic^52,53^. Thus, the mechanism and signals that link increased neuronal activity with glial lactate production must be independent from glutamate-induced glial glycolysis. One possibility is that NH_4_+ derived from active neurons, which is known to have an important role in the honeybee retina^54^, could act as the signaling molecule that stimulate glial lactate production. In this mechanism, neuronal mithochondria would produce NH_3_, which would be transformed to NH_4_+ given the external pH (6.7), then a transporter or a channel would allow the import to glial cells where it may stimulate pyruvate/lactate production by allosteric activation of phosphofructokinase^55^. Then, glial cells can export pyruvate/lactate or alanine to fuel neurons. Finally, alanine in neurons can be deaminated to produce ammonium. Although Volkenhoff and collaborators nicely demonstrated that glial cells in culture fueled by trehalose or glucose secrete alanine and lactate^14^, whether this mechanism is conserved *in vivo* remains to be established.

Several aspects of this system need to be determined. There are several putative genes that may act as *d*MCTs expressed in neural tissue that need to be confirmed in their function and importance, as well as their specific distribution in glial and neuronal cells. In addition, is important to stablish the role of MC transport in adult nervous system, more complex in the diversity of neurons and probably the energy need due to the more complex behaviors and longer life span of the adult fly. In summary, our work provides for the first time a description of monocarboxylate transport between glial cells and neurons from the ventral nerve cord of *Drosophila* larvae and reveals an increased lactate transport from glial cells to neurons during high neuronal activity demonstrating a role of Chaski mediating this lactate exchange.

## EXPERIMENTAL PROCEDURES

### Fly strains

Flies of either sex were raised at 25°C on standard *Drosophila* food composed by (per liter of media): 100 g yeast, 80 g glucose, 50 g wheat flower, 11 g agar, 6 ml propionic acid, 12 ml 20% Nipagin (Methylparaben). Flies used were: W^1118^ (experimental control), *OK6-Gal4*, *REPO-Gal4*, *R55B12Gal4* (#39103), *R83E12Gal4* (#40363), *R85G01Gal4* (#40436), *R54C07Gal4* (#50472), *AlrmGal4* (#67031) and *UAS-GCaMP6f* (#42747) from Bloomington Stock Center. *UAS-Pyronic* and *UAS-FLII^*12*^*Pglu700µ∆6 (glucose sensor)* flies were kindly donated by Drs. Pierre-Yves Plaçais and Thomas Preat (CNRS, Paris). UAS-*pHerry* was kindly provided by Dr Gregory MacLeod (Florida Atlantic University, Florida). *Chk*^*CRISPR*^* mutant was generated previously (Delgado, 2018), *UAS-Laconic* was generated in our lab and sent to inject by BestGene Inc. (California, USA).

### *Ex vivo* preparation of *Drosophila* VNC

Third instar larvae of each strain used were dissected in saline solution without Ca^2+^ containing (in mM): 128 NaCl, 2 KCl, 4 MgCl_2_, 5 trehalose, 5 HEPES and 35 sucrose, pH 6.7 or 7.2 as appropriate. The brain was removed without imaginal discs and placed on coverslip mounted on a recording chamber (Live Cell Instruments, Seoul) and perfused by a gravity-driven flow with saline solution composed by (in mM): 128 NaCl, 2 KCl, 1.5 CaCl_2_, 4 MgCl_2_, 5 trehalose, 5 HEPES and 35 sucrose at pH=6.7, except Supplementary figure 2, which was adjusted at pH 7.2, 6.7 and 6.0, at room temperature (23 °C). To correct for osmotic changes due to the different stimulations we used sucrose, which is not metabolized by *Drosophila*.

### Live cell imaging

Preparations were imaged in an Olympus BX61WI microscope equipped with DSU Spinning Disk Confocal System coupled to a DV2-emission splitting system (Photometrics, Tucson, USA) and a CCD camera Hamamatsu ORCA-*R*^2^ controlled by Cell^∧^R software (Olympus, Tokyo, Japan). Brains were visualized with an UMPlanFl 20X/0.5 water immersion objective. Images from ventral nerve cord (VNC) were for perineurial, subperineurial and cortical glial cells obtained from the most ventral part of the VNC; for the rest of cells analyzed we focused on the most dorsal part of the VNC (see figure 3A). For the FRET-based sensors excitation was 438/24 nm and emission were 488 nm (mTFP, CFP) and 540 nm (Venus, YFP) acquiring images every 30 seconds (1024×1024 pixels). For *pherry*, excitation was 488 nm for pHluorin and 540 nm for mCherry acquiring every 15 seconds. For *GCaMP6f* excitation was 488 nm, acquiring every 10 seconds.

To measure, we selected a region of interest (ROIs) in over ten cells for each brain using the software ImageJ (1.52e, National Institutes of Health, USA, depicted in figure 1A). The drift was corrected using the registration plug-in. Then, the mean grey value of each protein emission was measured and background-subtracted. The signal was the fluorescence ratio between mTFP/Venus (*Laconic* ratio or *Pyronic* ratio). To normalize, the mean value of two minutes before the stimulus of the mTFP/Venus ratio was used to divide each value after the stimulus unless otherwise were stated. For the glucose sensor, the ratio was YFP/CFP (*FLII^12^Pglu700µ∆6* ratio). For *pHerry* ratio, we divided the pHluorin signal to the mCherry signal. At the end of each experiment we exposed the brains to 5 mM of sodium azide as a positive control for FRET based sensors.

### pH and lactate levels in hemolymph

Hemolymph was collected of over ten W^1118^ larvae (68 hrs after larval hatching) washed three times in distilled water and dry in a paper towel. Then, a little incision was made with tweezers in the posterior part of the body of larvae and left to drain in a small concavity made in a parafilm. We measure lactate using 1 μL of the collected hemolymph using the enzymatic fluorometric assay (#K607, BioVision, CA, USA). Each determination was done by triplicate, in four independent experiments. For pH, the same protocol to collect the hemolymph was performed and measurements were done in the concavity made in the parafilm that contains the hemolymph drop with a microelectrode (Orion 9810BN, Thermo Scientific, IL, USA). We repeated the protocol in 20 different samples.

### Statistical analysis

All data were expressed as mean ± SEM. For statistical analysis, the GraphPad Prism 6 program was used. Multiple comparisons were performed with ANOVA followed by Bonferroni posttest. To determine if two sets of data were significantly different from each other, unpaired Student’s t-test was applied. p values less than 0.05 were taken as the significant limit.

---

**Supplementary figure 1.**
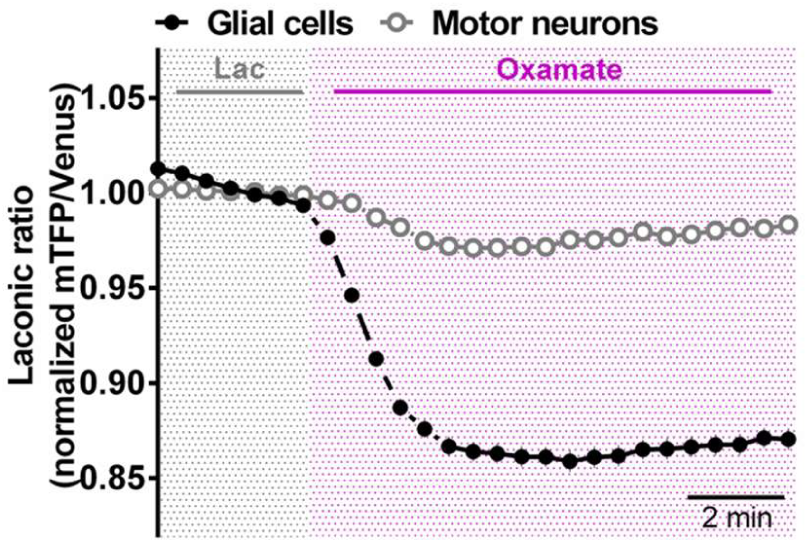
Trans-acceleration of *d*MCTs induced by oxamate shows lower basal levels of lactate in motor neurons than glial cells. Brains expressing *Laconic* in glial cells (*repoGal4>Laconic*, black circles) or motor neurons (*OK6Gal4>Laconic*, grey open circles) were incubated in saline solution with 1 mM lactate (Lac) and then depleted from lactate by exposition to 10 mM of oxamate. Values are the mean fluorescence ratio of mTFP/Venus obtained from over 50 cells from 5 independent experiments for each Gal4 used.

**Supplementary figure 2.**
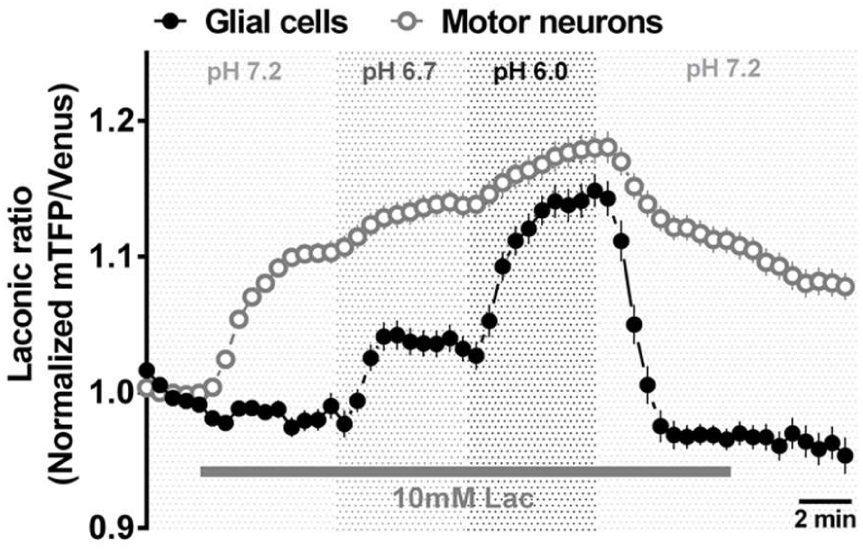
Monocarboxylate transport in *Drosophila* motor neurons and glial cells is dependent on extracellular pH. Brains expressing *Laconic* in all glial cells (*repoGal4>Laconic*, black circles) or motor neurons (*OK6Gal4>Laconic*, grey circles) were bathed in saline solution at pH=7.2 and imaged every 30 seconds. Then brains were exposed to 10 mM of lactate and to a decrease in pH levels (ranging from 7.2 to 6.0). Mean fluorescence ratio of mTFP/Venus (*Laconic*) was obtained from over 50 cells from 4 independent experiments.

